# Epidermal stem cell compartment remains unaffected through aging in naked mole-rats

**DOI:** 10.1101/2020.11.13.381061

**Authors:** A. Savina, T. Jaffredo, F. Saldmann, C.G. Faulkes, P. Moguelet, C. Leroy, D. Del Marmol, P. Codogno, L. Foucher, M. Viltard, G. Friedlander, S. Aractingi, R.H. Fontaine

## Abstract

Skin represents an informative and convenient organ for the analysis of the aging process. Naked mole-rats (NMR) are subterranean rodents remarkable for their longevity, with unexplained resistance to skin aging. In middle-aged NMR, extensive *in situ* analysis indicated that skin compartments and cell types remained similar to young animals. Using single-cell RNA-sequencing, we found three classical cellular states defining a unique keratinocyte differentiation trajectory that did not appear to be altered during aging after pseudotemporal reconstruction. Finally, NMR skin healing closure was strictly comparable between the two age groups. These results indicate that the content in stem cell populations as well as the differentiation process are preserved during aging in NMR and that such properties are related to the healing process.

## INTRODUCTION

Naked mole-rats (NMR, Heterocephalus glaber) are small poikilothermic and hairless rodents native to East Africa, where they live underground in eusocial colonies. Although displaying a similar size than the mouse, NMR live almost ten times longer with a maximum lifespan exceeding 30 years in captivity. However, in their natural habitat due to increased mortality risks, NMR live far shorter lifespans - commonly up to around 10 years, although occasionally more than 17 years(*1*–*3*). Despite being the longest lived rodent, NMR do not show any increase in age-specific hazard of mortality in defiance of Gompertz’s law(*4*) and all of the normal signs of aging such as decrease of fertility, muscle atrophy, bone loss, changes in body composition or metabolism seem to be mostly absent in these animals(*5*). Moreover the incidence of age-related diseases such as metabolic disorders, neurodegenerative diseases and even cancers are still absent or extremely low in NMR (*6*, *7*). The NMR genome has been sequenced in 2011 revealing a unique genetic profile and molecular adaptations correlated with all the characteristics described above: resistance to cancers, poikilothermia, absence of pelage, insensitivity to low levels of oxygen, as well as impaired vision, taste and circadian rhythms. 93% of the NMR genome has a synteny with the human, rat or mouse genome(*8*). Thereby, NMR emerges as an important non-model organism, with high informative value in research for the study of aging.

To date, many of the NMR anti-aging mechanisms are poorly understood. Further, only two descriptive scientific articles have focused their research on NMR’s skin investigating how the skin morphology of these animals is adapted to thermoregulation and their subterranean environment(*9*, *10*). However, skin remains an ideal organ for the analysis of several types of biological phenomena that can be associated with aging. Skin is divided into three main layers: epidermis, dermis and hypodermis. The outermost layer, the epidermis, is composed itself of four tightly adherent layers: basal, spinous, granular and corneous. Following injury, a wound healing response is rapidly activated to repair and restore the skin’s barrier function. Both homeostasis and repair of the epidermis are maintained by specific keratinocytes stem cells or progenitors found in the basal layer, proliferating and providing new cells for the upper suprabasal differentiating layers(*11*). Aging in rodents and humans is attributed to loss or lineage-skewing of these resident stem cells, altering normal homeostasis and tissue *repair(12–14*). In order to study the potential effects of aging on NMR epidermis and the putative role of stem cells, we used single-cell RNA-sequencing (scRNA-seq) to decipher the molecular characteristics of young and middle-aged NMR epidermal cell populations and also analyzed in depth their epidermal morphology. Finally, since repair is one of the main skin duties that alters lifelong, we analyzed skin healing in NMR during aging.

## RESULTS

### ScRNA-seq of NMR epidermis defines 3 cellular states for NMR keratinocytes

First, to study the global transcriptional heterogeneity of the NMR epidermis using scRNA-seq, dorsal skin of 3 young (Y, mean age: 1.1 years) and 3 middle-aged (A, mean age: 11.3 years) NMR were collected. After a short enzymatic epidermis-dermis digestion and separation, epidermal cells were mechanically dissociated and directly loaded into the chromium controller (10x Genomics) without any FACS-sorting to prevent cells from stress (Fig.1A). All the samples were processed in parallel to minimize batch effects.

**Fig.1.**
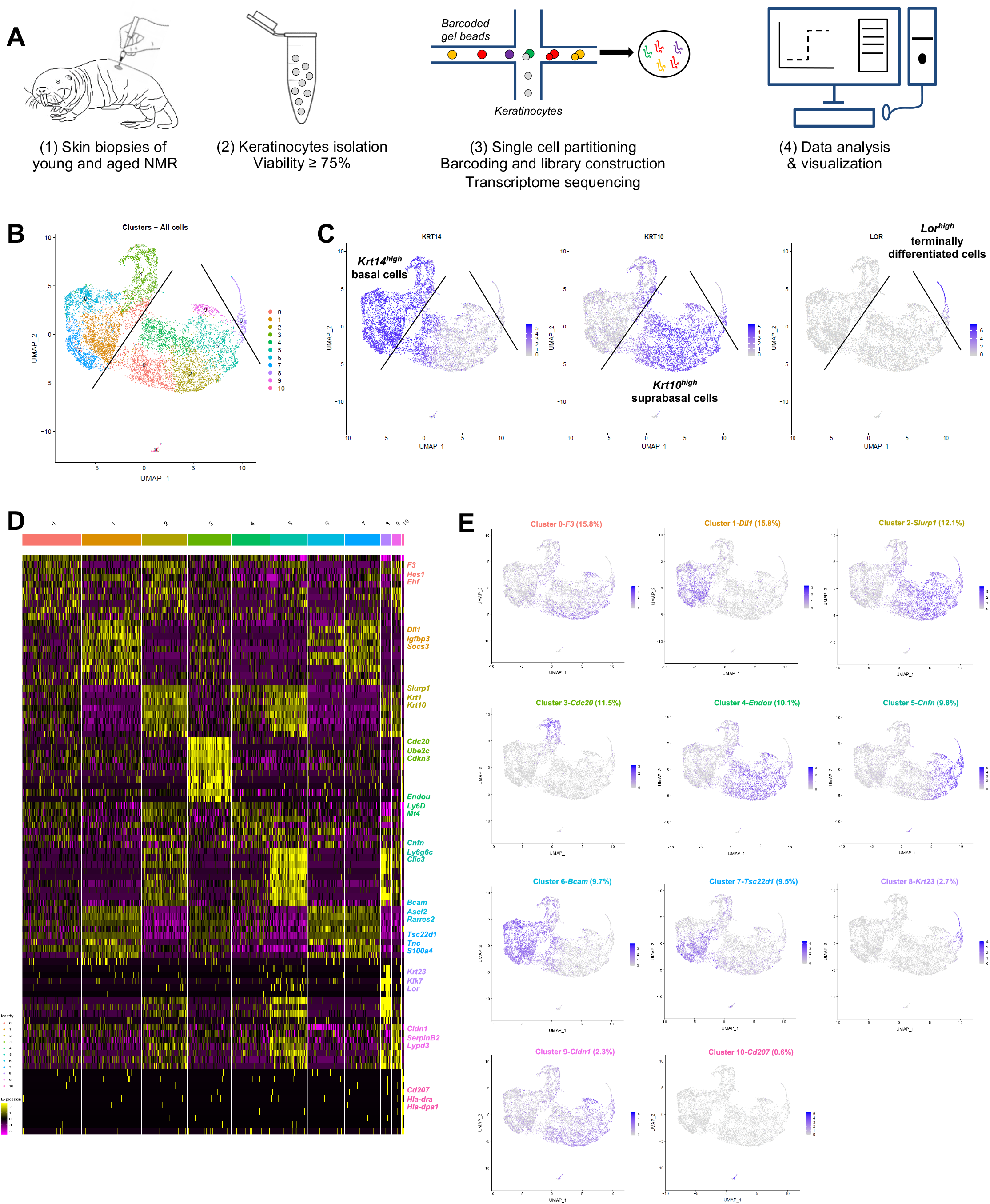
Cell populations characterization of naked mole-rat (NMR) epidermis using single-cell RNA-sequencing (ScRNA-seq). (**A**) Overview of the experimental workflow. (**B**) Global epidermal cell transcriptomes (n=10,232 cells) visualized with Uniform Manifold Approximation and Projection (UMAP), colored according to unsupervised clustering to better compare different cell subtypes. 11 different cell clusters were found, numbered from 0 to 10. (**C**) Expression of the 3 main marker genes during epidermal differentiation *(Krt14, Krt10, Lor)* projected onto UMAP. Cells were sub-divided into 3 main groups according to the differential epidermal cellular states: basal *(Krt14^high^),* suprabasal/intermediate *(Krt10^high^)* and corneous *(Lor)* layers. Cells with the highest expression level are colored dark blue. (**D**) Heatmap of differentially expressed genes. For each cluster, the most differentially expressed genes and their relative expression levels in all sequenced epidermal cells are shown. Cells are represented in columns, and genes in rows. 3 selected genes for each cluster were color-coded and annotated on the right. (**E**) Expression levels for each cell are color-coded and overlaid onto UMAP plot for one selected cluster-specific gene. Cells with the highest expression level are colored dark blue. % of cells in each cluster is annotated.

Quality control (# UMI ≥ 2000, # genes ≥ 700, % MT ≤ 25%) made with FastQC, Cellranger and Seurat v3 pipeline (Fig. S1) for unsupervised clustering revealed that one sample from the A group had a very distinctive pattern after Principal Component Analysis (PCA). This sample was removed, keeping for the rest of our study 3 samples in the Y NMR group and 2 samples in the A NMR group (see materials and methods). In total, we obtained 10,232 cells from 5 NMR epidermis for downstream analyses.

Based on gene expression profiles and Uniform Manifold Approximation and Projection (UMAP) to visualize cell clusters, we identified 11 cellular populations (numbered from 0 to 10, Fig. 1B) and annotated them with differential expression testing and a list of 98 selected genes of interest for skin homeostasis and repair(*15*, *16*). We found 2 different populations: keratinocytes representing 99,4% (10,171 cells) in clusters 0 to 9, and immune cells in cluster 10 representing 0.6% of epidermal cells (61 cells). This immune isolated cluster 10 contained a majority of *Cd207^+^/Ctss^+^/Mfge8^+^* Langerhans cells(*17*, *18*).

Cytokeratin expression can globally define the differentiation status of keratinocytes: Keratin 5 and 14 *(Krt5 and Krt14)* are expressed in active basal cells, while Keratin 1 and 10 *(Krt1 and Krt10)* are found in differentiating suprabasal cells and Loricrin *(Lor)* occupies 80% of the epidermal corneous envelope(*19–21*). Thus, using these well-defined marker genes, we were able to confirm that here too, the keratinocyte population separates into 3 main cellular states. Four clusters (clusters 1-3-6-7) expressed high levels of *Krt14 (Krt14^high^)* and *Krt5* and were classified as basal cells. Clusters 0-2-4-5-9 expressed higher levels of *Krt10 (Krt10^high^)* and *Krt1* and were categorized as suprabasal or intermediate cells. The last keratinocyte-cluster (cluster 8) was the only one to express very high levels of Loricrin *(Lor^high^)* and was considered as a cluster of terminally differentiated keratinized cells of the corneous layer (Fig. 1C). The same results were confirmed with violin plots of these marker genes (Fig. S2A) and t-distributed Stochastic Neighbor Embedding (t-SNE) visualization (Fig. S2B-C). The most differentially expressed marker genes for each cluster were represented using heatmap visualization (Fig. 1D) and 3 specific genes for each cluster were highlighted and color-coded to match cluster color. For each cluster, the most representative gene was projected onto the UMAP plot (Fig. 1E).

In order to confirm the 3 cellular states in NMR keratinocytes, we identified other well-known skin layer genes in each cluster. In *Krt14^high^* basal cells clusters, cells from clusters 1-6-7 expressed specific basal layer genes such as *Tp73, Itga6, Itgb1, Cxcl14 and Smoc2*(*15, 21–23*). Cells from cluster 1 expressed the Notch ligand *Dll1,* a marker of stem cells found in the basal layer in human and mouse skin(*15*, *24*), *Igfbp3*, a gene exclusively expressed in the basal layer’s proliferative keratinocytes in normal skin(*13*, *15*, *25*, *26*), and *Socs3,* a gene known to promote maintenance of epidermal homeostasis(*15*). Of note, *Dll1* and *Igfbp3* were also expressed in the two other basal clusters 6 and 7. Cluster 6 cells expressed *Bcam, Ascl2* and *Rarres2,* 3 genes found in the basal layer of the epidermis and known to stimulate keratinization(*27*, *28*). *Tsc22d1, Tnc, S100* family members (such as *S100a4* and *S100p)* and *Ccl2* expressed in Cluster 7 cells were reported to be restricted to basal/follicular stem cells and progenitors with high cell turnover notably during wound healing(*15*, *26*, *29*)*. Krt14^high^* cluster 3 cells specifically expressed classical epidermal cell cycle regulators such as *Cdc20, Ube2C and Cdkn3(26, 30, 31)* (Fig. 1E). Concerning *Krt10^high^* suprabasal cells clusters 0-2-4-5-9, we found that *Notch* receptor gene was expressed in clusters 5 and 9 cells, while its target gene *Hes1* was found in cluster 0 and 4. It has been demonstrated that *Notch* activation induces terminal epidermal differentiation through *Hes1* which is expressed in the spinous layer. The expression of another Notch ligand, *Jag2* was restricted to basal layer’s clusters 1-6-7 cells corroborating previous findings. *Jag1* on the other hand was only found in suprabasal cluster 9 cells(*24*). In addition to *Hes1,* cluster 0 cells also expressed *F3* and *Ehf* reported to be located in suprabasal layers of the skin playing an essential role in keratinocyte differentiation *15*, *32*). Cluster 4 analysis revealed the expression of *Endou, Ly6D* and *Mt4,* 3 genes previously associated with keratinocyte differentiation in the spinous layer of the skin(*15*, *23*). Of note, *Mt4*, reported to specify a transitory stage from basal to spinous layer(*15*, *29*), was expressed in cells from cluster 0, 4 and 2. Cluster 2 cells strongly showed classical suprabasal gene expression such as *Krt1, Krt10* and *Slurp1,* known to be a marker of late differentiation state, predominantly expressed in the granular layer of the skin(*33*). In cluster 5 cells, *Cnfn, Ly6g6c and Clic3* expression matched with a granular layer phenotype(*15*). The last suprabasal cluster defined here, cluster 9, was also consistent with genes expressed by differentiating keratinocytes in the granular layer of human and mouse skins such as *Cldn1, SerpinB12* and *Lypd3(15*). In addition, both clusters 5 and 9 cells revealed the expression of several strong makers such as *Hspb1* and *Grhl1* suggesting the possible granular keratinocytes phenotype of these two cluster cells(*15*, *31*) (Fig. 1E). Finally, in addition to *Lor,* cluster 8 cells expressed *Klk5/7, Krt23* or *Cdsn* strongly suggesting these cells were terminally differentiated keratinocytes from the corneous layer (*15*) (Fig. 1E).

The single cell transcriptomic analysis of NMR keratinocyte unraveled, for the first time in NMR, an epidermal differentiation process that follows 3 classical cellular states, from basal layer’s proliferative stem cells and/or progenitors (clusters 1-3-6-7) to suprabasal (clusters 0-2-4-5-9) and terminally differentiated cells (cluster 8). Within each of these 3 cell population, each cluster could represent a specialized subpopulation.

### Cell populations and trajectories analysis of NMR keratinocytes reveals no differences between young and middle-aged animals

Then, to study the transcriptional changes of the NMR epidermis during aging, we compared the Y and A NMR epidermis. Three Y NMR of 1.1 years old (n=6,014 cells) and two A NMR of 11.3 years-old (n=4,218 cells) were analyzed. Young and middle-aged cells seemed to be equally spread across clusters using UMAP visualization (Fig. 2A and 2B). This observation was also retrieved using t-SNE visualization (Fig. S3A). In addition, the clusters’ cellular states, defined by the expression of *Krt14, Krt10* and *Lor* marker genes and visualized with violin plots, remained the same between A and Y animals (Fig. 2C). The percentage of cell types between the 2 conditions after normalization was not statistically significant in every cluster (Chi^2^ statistic test pval ≤ 0,05; Fig. 2D). To identify differentially expressed genes between Y and A animals, we then compared the expression of genes between all cells from Y samples with all cells from A samples. No gene was found differentially expressed using the following thresholds: p-value below 0.05 and average log fold-change (logFc) above 1.

**Fig. 2.**
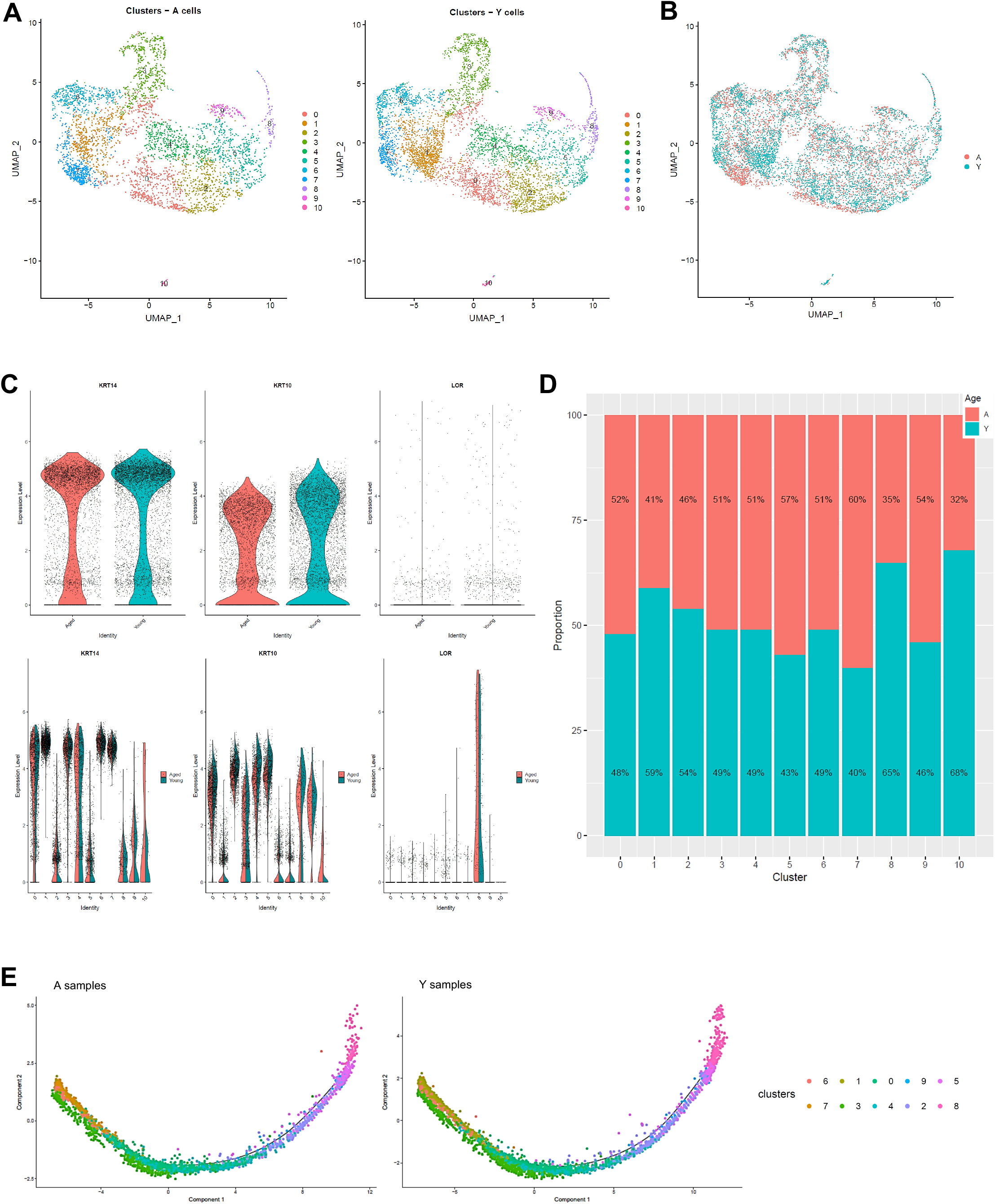
Single cell populations and trajectories analysis of young versus middle-aged NMR epidermal cells reveals no difference between the 2 age groups. (**A**) Uniform Manifold Approximation and Projection (UMAP), colored according to unsupervised clustering from middle-aged (A; n=4,218 cells) and young (Y; n=6,014 cells) NMR epidermal cells. Cell populations from both A and Y NMR were unchanged as the same 11 clusters were detected. (**B**) Cells from both A (red) and Y (blue) NMR were jointly projected on the same UMAP plot, showing an overlap of the 2 age groups tested. (**C**) Violin plots of *Krt14, Krt10* and *Lor* marker genes expressed by A (red) and Y (blue) epidermal cells (upper panel) and segregated by cluster (lower panel). No detectable changes in their expression were found. (**D**) Bar graph representing the relative proportion of epidermal cells in each cluster between A (red) and Y (blue) animals. There was no significant difference in epidermal cell proportion between the 2 age groups using Chi^2^ statistic test (pval ≤ 0,05). (**E**) Unsupervised differentiation trajectories constructed with M3Drop and Monocle v2.10.1 for A and Y keratinocytes. Immune cells’ cluster 10 has been excluded to focus on epidermal keratinocytes. The results showed a unique trajectory with no branches in both samples and the same repartition of cluster among the trajectory.

However, as subtle differences might exist, a deeper analysis between Y and A animals was performed separately within clusters. Using a p-value below 0.05 and an average log fold-change above 1, only 2 differentially expressed genes for basal cluster 7 *(ENSHGLG00000002542/Rps2* and *Igfbp3)* and 5 differentially expressed genes for immune cluster 10 *(S100a9, ENSHGLG00000004445/S100a8, Krt17, S100p,* and *Krt5)* exceeds these thresholds and were overexpressed in A animals. No differentially expressed genes typical of Langerhans cells were found. Conversely, in mice and humans, skin aging has been associated with reduced *Igfbp3* expression, whose localization in the basal/germinative layer of the epidermis suggests a key role in modulating epidermal homeostasis(*13*, *15*, *25*, *26*). S100 proteins, especially S100A8/A9 who exert anti-inflammatory function in healthy state, and Keratin intermediate filament proteins have been proved to be novel important regulators of inflammation and immunity in skin.

As keratinocytes differentiate, they undergo a process of transcriptional reconfiguration. Using Monocle 2, we reconstructed the single cell trajectory of Y and A NMR keratinocytes to mimic their biological differentiation process. Of note, cluster 10 cells corresponding to epidermal immune cells was excluded from all Monocle analysis to focus on the keratinocytes’ differentiation program. Colored by cluster, we found the same linear trajectory for each age condition, with no branches, suggesting an absence of different cellular decision or alternative expression program between Y and A keratinocytes (Fig. 2E). For both Y and A NMR, the distribution of keratinocytes along their process of cellular differentiation showed that, as expected, *Krt14^high^* basal clusters 1-3-6-7 cells were condensed at the root of the trajectory process, with cluster 3 cells spreading out more from this root showing their early engagement in the differentiation process. Then, *Krt10^high^* intermediate clusters 0-2-4-5-9 cells moved towards the right of the trajectory, as they were more committed in the differentiation process (no cells were observable in the root). And finally, *Lor^high^* cluster 8 cells were distributed at the very end of the trajectory, for terminally differentiated keratinocytes(*34*) (Fig. 2E). These results were also represented by Monocle states (state or “segment of the tree”, Fig. S3B) and for each cluster separately (on separate plots) for merged samples or Y and A samples separately, to see precisely where each cluster was located Fig. S3C).

Colored by pseudotime, the cells were also visualized according to a blue gradient which becomes lighter as cells move away from the root of the trajectory, reflecting their maturation state. No significant difference on pseudotime estimation was found between Y and A NMR (Fig. 3A). To validate our 3 cellular states in NMR keratinocytes, the 3 selected well-known gene markers of epidermal differentiation *Krt14, Krt10* and *Lor(34)* were plotted according their mean expression levels (black line) along the pseudotime axis (Fig. 3B). No changes between our 2 age groups was noted suggesting an absence of age-related effect on keratinocyte differentiation in NMR (Fig. 3B). To assign a more precise cellular state of differentiation to the 10 clusters, we chose to individualize them on separate plots (Fig. 3C). Thus, combined with data relative to specific gene expression obtained in Fig. 1, pseudotime reconstruction allowed us to order the 10 clusters precisely and to partition them into 5 metaclusters (i.e group of clusters) according to their differentiation engagement: i) basal 1 containing clusters 6-7-1 corresponding to basal keratinocytes stem cells / progenitors; ii) basal 2 for cluster 3 only corresponding to mitotic or proliferative stem cells; iii) intermediate 1 containing clusters 0-4 and comprising cells characteristic of spinous layer cells, iv) intermediate 2 aggregating clusters 2-5-9 containing granular layer cells and finally, v) terminally differentiated cells named corneum for cluster 8 (Fig. 1, 3C). The expression of *Krt14, Krt10 and Lor* marker genes visualized with violin plots confirmed the cellular states of these newly defined metaclusters Fig. S4A). The repartition of cell types between the Y and A animals after normalization was not statistically significant in each metacluster (Chi2 statistic test pval ≤ 0,05; Fig. S4B). Distributions of the frequency of Y and A cells as a function of the pseudotime on its scale were also plotted. Wilcoxon test revealed that Y and A NMR keratinocytes distributions was similar using the pseudotime estimation (p-value = 0.1673). According to pseudotime segments, no differential gene markers were found between the segments (Fig. S4C). Then, we observed the most differentially expressed genes and their relative expression levels in all sequenced epidermal cells on a heatmap (Fig. 3D). Similar to the previous analysis, five gene metaclusters could be clearly identified, each with a different region of activity during the differentiation process. Unsupervised differentiation trajectories for Y and A animals showed identical repartition of these metaclusters among the trajectories (Fig. 3E).

**Fig. 3.**
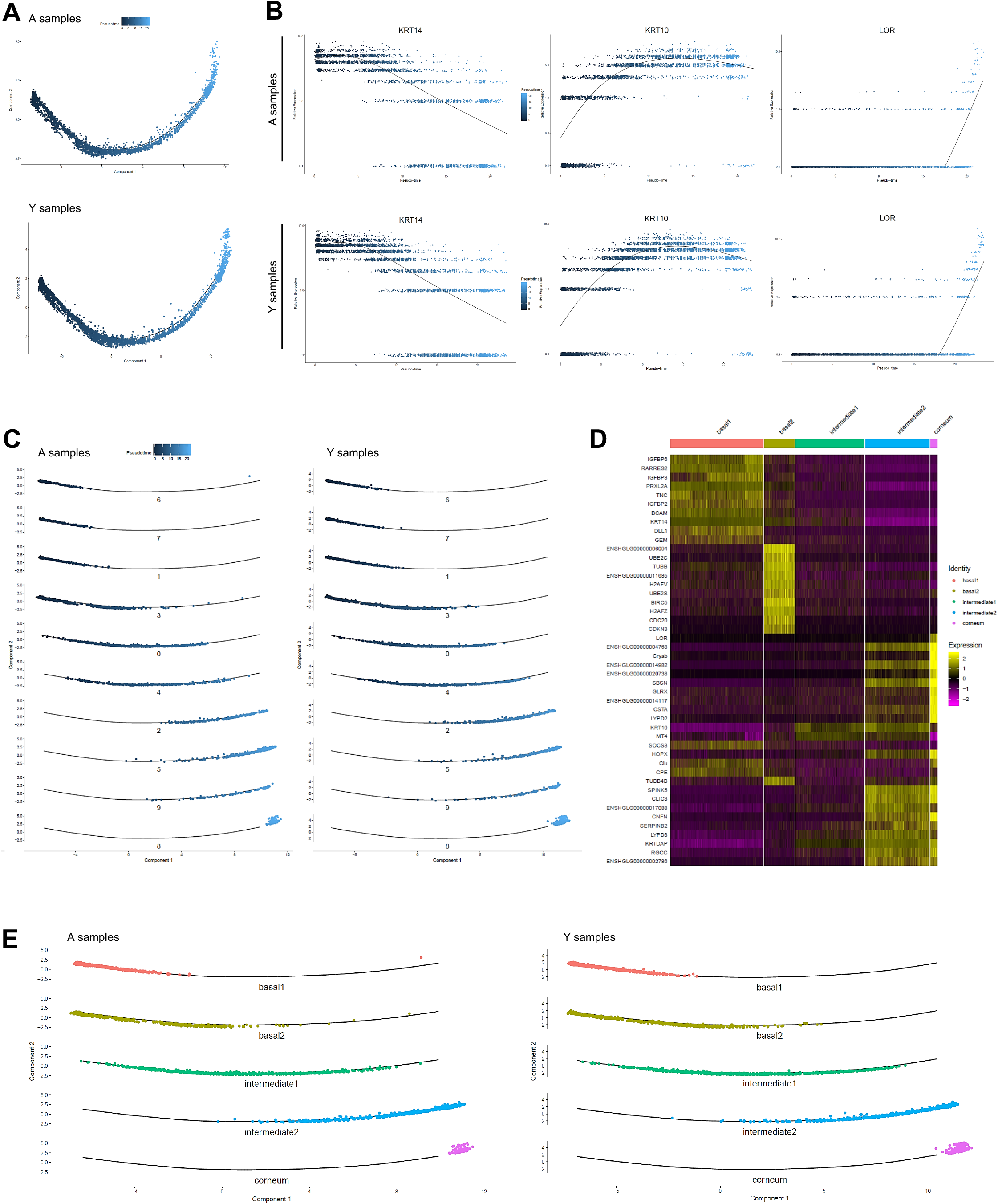
Pseudotime analysis and metacluster identification of young versus middle-aged NMR keratinocytes. (**A**) Colored pseudotemporal ordering of keratinocytes for A (middle-aged) and Y (young) NMR separately. No significant differences were observed between the 2 groups. (**B)** Relative expression of *Krt14, Krt10* and *Lor* markers plotted along the pseudotime axis for each condition. The same relative expression profile was found between A and Y NMR, validating the 3 cellular states for NMR keratinocytes and the no age-related impact of aging in this keratinocyte differentiation process. (**C)** Pseudotime estimation showing epidermal progression along the differentiation process with individualized cluster visualization, for each condition. Five metaclusters were defined according to their differentiation engagement: basal 1, basal 2, intermediate 1, intermediate 2 and corneum. (**D**) Heatmap of most differentially expressed genes for each newly defined metacluster. Cells are represented in columns, and genes in rows. (**E**) Unsupervised differentiation trajectories for A and Y animals. The same repartition of metacluster among the trajectory was observed.

To conclude, we observed 11 equivalent clusters defining epidermal cells from Y and A NMR and a single path trajectory. Using pseudotemporal ordering, we were able to identify 5 metaclusters. Basal metaclusters containing keratinocytes stem cells and proliferative progenitors seemed to contribute to all the other cellular states to promote epidermal differentiation, up to the upper corneous layer. In all of these results, by comparing each time Y and A NMR keratinocytes, we did not observe any age-related effect, contrary to what has been described in mouse or human(*13*, *21*).

### Metacluster characterization and enrichment analysis revealed specific functions for NMR keratinocytes

As basal cells transit through different cellular states until terminal differentiation, we examined the dynamics on the genes used to construct the trajectories using Monocle. We identified 6 gene modules that followed similar kinetic gene changes along pseudotime. The heatmap profiles were identical, suggesting that the same gene groups at the same expression level were orchestrating the differentiation process in both Y and A NMR keratinocytes, independently of aging (Fig. 4A, left panel). We then investigated whether specific functions could be assigned to those 6 groups using Gene Ontology (GO). The most significant GO terms found in each activated gene group for both ages comforted their implication in their commitment : early activated cluster 5 for basal layer state, cluster 1 for cell cycle still in the early pseudo-timeline (note the sharp transition between these 2 states), mid-time activated cluster 6 for cell differentiation, cluster 3 at the end of differentiation program for an ultimate transition, cluster 2 strictly circumscribe at the end of pseudotime differentiation process and finally cluster 4 harbouring a biphasic expression pattern with functions involved at various time of the differentiation process supported by different genes (Fig. 4A, right panel).

**Fig. 4.**
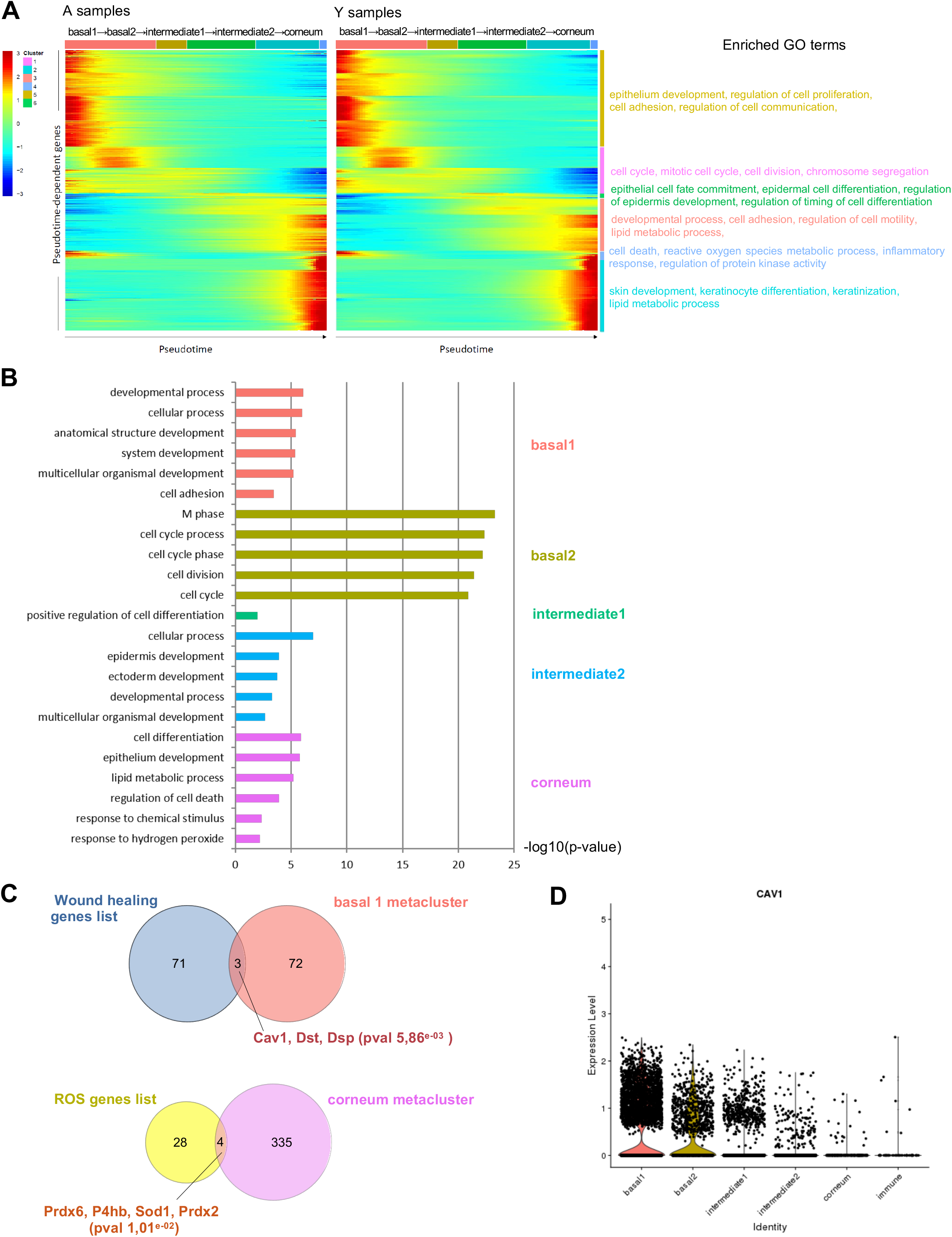
Metacluster characterization of NMR keratinocytes. (**A**) Heatmap pf pseudotemporal dynamics of the pseudotime-dependent genes in A (middle-aged) and Y (young) NMR (left panel). Each row (i.e., gene) is normalized to its peak value along the pseudotime. 6 distinct clusters or stages during pseudotime are represented by colored bars on the side. The most significant enriched GO terms in each gene cluster are listed (right panel). The gene ordering is strictly the same between A and Y heatmaps. (**B**) Barplot representing a selection of the most significant enriched Gene Ontology (GO) terms (from BiNGO analysis) in each keratinocytes metaclusters (except immune cluster) sorted by p-value. Barplot represents the adjusted p-value. (**C**) Venn diagram depicting the genes overlap significantly overrepresented in basal 1 ((pval 5,86^e-03^) and corneum metaclusters (pval 1,01^e-02^) as compared respectively with wound healing and ROS gene lists. (**D**) Violin plot representing *Cav1* gene expression enrichment in metaclusters. Note that its expression is the strongest in basal 1 metacluster.

The analysis between Y and A animals within metaclusters with a p-value below 0.05 and an average logFc above 1 revealed only for immune cluster 10 the same 5 differentially expressed genes *(S100a9, ENSHGLG00000004445/S100a8, Krt17, S100p,* and *Krt5*). No statistical differences were observed using these thresholds for the other metaclusters.

Next, another GO analysis was performed on the different epidermal metaclusters defined above. We used the most representative markers of each metacluster and described a repartition of function for each of them (Fig. 4B). Basal 1 metacluster (cluster 1-6-7) was characterized by developmental and cell adhesion processes relative to their basal stem cell/progenitor phenotypes. Indeed, those cells were likely prone to establish cell junction to extracellular matrix (ECM) and to commit to a global developmental process of tissue formation. As expected, metacluster basal 2 (cluster 3) was well defined by a wild range of cell cycle functions relative to their mitotically active progenitors cell type. Intermediate 1 (cluster 0-4) and Intermediate 2 (2-5-9) metaclusters were characterised by cell differentiation and developmental processes confirming these clusters as a population of differentiated keratinocytes of the spinous and granular layers respectively. Lastly, Corneum metacluster (cluster 8) was, as expected, represented by active lipid metabolism as well as cell death regulation and reactive oxygen species (ROS) reaction terms (Fig. 4B). The characteristics of metaclusters were well maintained in both age groups. The global functional repertoire of this newly sequenced transcriptomes described here could also be compared to one or more existing transcriptomes of related species.

Furthermore, based on gene lists associated to synonymous terms, we performed an enrichment analysis for our metaclusters for aging, ROS, and wound healing. Surprisingly, only a few genes were significantly enriched for ROS and wound healing lists; no enrichment was retrieved from the aging gene list (Fig. 4C). Corneum metacluster (cluster 8) was enriched in 4 genes related to ROS function: *Prdx6, P4hb, Sod1* and *Prdx2* (Fig. 4C; pval 1,01^e-02^). In NMR, the abundantly expressed antioxidant enzyme *Sod1* activity was 1.35-fold higher as compared to age matched mice but age did not have an effect on its activity in both species. Peroxiredoxins, especially *Prdx2,* required for extended lifespan at lower temperature, were shown to be expressed in NMR livers, which may result in an increased levels of ROS and suggest that the long-lived NMR can thrive despite elevated oxidative stress(*35*).Of note, *Prdx 2* and *5* were also enriched in the immune cluster. Those immune cells expressed high levels of transcripts related to regulation of immune system process and response to stimulus, showing that their main function is to be involved in resistance to skin pathogens. Those terms were also found at a lower level in the corneum metacluster where immunity meets external antigens and in the basal 1 metacluster where keratinocytes have key roles in immune defense Fig. S5). As the contributions of epidermal stem cells in wound healing and tissue regeneration have been extensively studied in mouse and humans(*26*, *36*) but never in NMR, we decided to focus our attention on wound healing. In basal 1 metacluster, 3 genes were significantly enriched from a list of 74 genes: *Cav1, Dst, Dsp* (Fig. 4C; pval 5,86^e-03^). Mutations in the mouse *dystonin (Dst)* and *desmoplakin (Dsp)* genes have previously been shown to cause skin fragility(*37*, *38*). *Caveolin 1 (Cav1)* was strongly expressed in each basal cluster (6, 7 or 1) of the basal 1 stem cells/progenitors metacluster (Fig. 4D). *Cav1* has been found in undifferentiated epidermal basal layer keratinocytes and considered by others as a potential epidermal stem cell markers(*39*). Moreover, some studies have proven that *Cav1* could be crucial as a promoter of the wound healing process. Overexpression of *Cav1* in epidermal stem cells promoted their proliferation and re-epithelialization, a necessary stage for restoring the epidermis during wound healing(*40*).

### Histomorphology of young versus middle-aged NMR skin

To further investigate skin physiology and healing during aging in these animals, we performed classical staining and immunohistochemistry on the skin of Y and A NMR. HE and Sirius Red stainings allowed us to measure epidermal and dermal thickness of back skin samples (Fig. 5A-B). Surprisingly, A NMR showed a global histomorphological structure similar to that of the Y NMR, but the epidermis of the A animals was significantly thicker. No effect of age was observed on the dermal thickness (Fig. 5C). The number of epidermal cells Fig. S6A) and epidermal layers Fig. S6B) were significantly higher in the A group. More precisely, cell counting per layer revealed an additional cellular layer (“layer 6”) in older animals Fig. S6C). Masson’s trichrome staining for collagenous connective tissue fibers revealed no difference between the 2 groups just like hyaluronic acid, a linear polysaccharide of the extracellular matrix contributing to contact inhibition and cell proliferation(*41*) (Fig. 5 D-E). Classical epidermal layers antibodies such as anti-Keratin14 (basal layer marker), Keratin10 (suprabasal layer marker), Loricrin (a major protein of the cell corneous envelope), Filaggrin (intermediate filament-associated protein of the outer granular layer of the epidermis), Laminin 5 (dermal-epidermal basement membrane protein), Integrin-beta4 (ITGB4, basal surface marker juxtaposed to the dermal-epidermal basement membrane) or collagen I (found in all dermal layers) showed no difference between the 2 age groups (Fig. 5F-L). It has been discovered that epidermal proliferation declined with age especially in humans(*13*). Interestingly, in NMR, Ki67 index for cell proliferation revealed no significant difference between Y and A NMR (Fig. 5M). Finally, p16, one of the most frequently used markers of senescence and an important hallmark of ageing, was detected in NMR skin but no differences were observed between the 2 groups (Fig. 5N) whereas this marker accumulate in an age-dependent manner in mammalian skin(*42*).

**Fig. 5.**
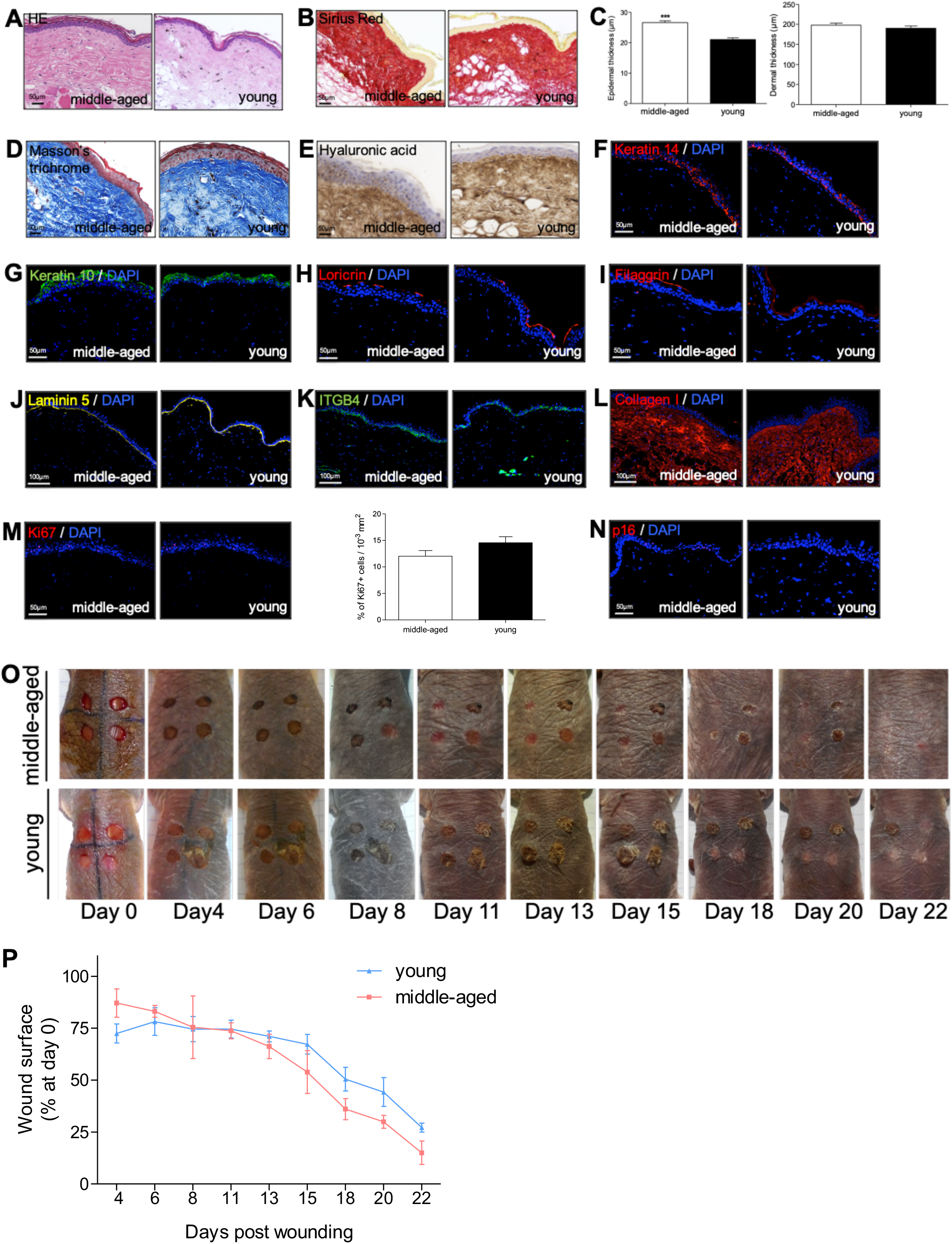
NMR skin histomorphology and wound healing. (**A, B, D**) HE, Sirius Red and Masson’s Trichrome staining photomicrographs on sections of middle-aged (A) and young (Y) NMR back skin. (**C**) Histograms showing the epidermal and dermal thickness (μm) of middle-aged and young NMR back skin. ***p≤0.0001 in Mann-Whitney U test. n=4 animals per group. (**E-N**) Hyaluronic acid, Keratin 14, Keratin 10, Loricrin, Filaggrin, Laminin 5, ITGB4, Collagen I, Ki67, and p16 photomicrographs on sections of middle-aged and young NMR back skin. Cells were stained with either Alexa 546 or Alexa 488 fluorochromes. For hyaluronic acid, cells were stained with 3,3’-diaminobenzidine. Negative controls were performed in parallel with the samples substituting the primary antibody with the equivalent isotype. Histogram showed the number of Ki67+ cells per surface in both age group. Scale bar = 50 or 100 μm. *represents differences between A and Y animals. Bars: SEM. ***p≤0.0001 in Mann-Whitney U test. n=4 animals per group. (**O**) Photomicrographs of back skin wounds for middle-aged and young animals from day 0 to day 22 post wounding. (**P**) Wound closure rate (% of wound surface at day 0) from day 4 to day 22 post wounding. No statistical difference was found the 2 groups using Mann-Whitney U test. n=4 animals per group.

### Wound healing is not affected by age in NMR

The capacity of skin to heal quickly and completely is a major function that features the skin status. Aging could also play a major role in altering wound healing(*43*). We therefore compared skin healing in Y and A NMR. Punch biopsy devices were used to create surgical wounds on the backs of Y and A NMR and the wound closure rate was measured (percentage of wound closure at day 0) for the first time in these animals. Unlike normal mice that possess a healing time course of approximately 5 days, NMR appeared to heal at a slower rate of 22 days and with less scarring (Fig. 5O). Wound closure has also been proved to occur more rapidly in young than in aged mice, or humans(*14*). Here, in NMR, the wound closure rate of A animals followed the same progression than the one of Y animals (Fig. 5P).

## DISCUSSION

Age-related epidermal changes rely essentially on a reduced keratinocyte stem cell compartment, reduced proliferation, reduced epidermal thickness, increase in their turnover time and flattening of the dermal-epidermal junction. Here, using molecular and cellular experiments, we demonstrated that NMR epidermal homeostasis remained unaltered for the first 11 years of their life. This age would represent an old animal in the wild where few animals live beyond 10 years and most die at age 1-2 years, and a middleaged individual for most captive animals where mortality increases rapidly beyond 25 years(*1–3*).

The epidermis is composed of 3 major cell types: keratinocytes, melanocytes and immune cells, notably Langerhans cells. In NMR epidermis, we did not find cells expressing melanocytes genes, which is consistent with previous reports showing that melanocytes were only detected in the NMR dermis(*9*). Thus, NMR epidermal cells displayed 2 populations: keratinocytes and immune cells. By ordering our 10 keratinocytes clusters, we found a “classic” differentiation process including basal stem cells and proliferative keratinocytes, followed by spinous then granular layer cells and finally terminally differentiated cornified cells in both young and middle-aged NMR. Interestingly, analysis of basal keratinocytes stem cells cluster 7 in young versus middle-aged animals revealed a threefold increased *Igfbp3* expression in older animals. Conversely, in mice and in humans there is a large decrease in *Igfbp3* during skin aging(*13*). Thus, we hypothesize that maintenance of high basal *Igfbp3* could be one of the mechanisms allowing the cutaneous resistance to aging in NMR skin.

Even if age-related changes in skin stem cells homeostasis still remain under debate, our results are different from what has been previously shown in other rodent species and humans(*12*, *13, 21, 31*). Indeed, in NMR, the same number of epidermal clusters and metaclusters harbouring equivalent functions, cellular states and differentiation trajectories were conserved at older age. Of note, it has been recently shown that individual cell clustering analysis was indeed a powerful tool to detect differences between young and adult mice epidermis since young epidermal cells progenitors appeared as a more homogenous compared with adult cells progenitors(*26*). Moreover, these authors found only one differentiation path in young mice but 2 differentiation paths in older mice, while others have even found 3 distinct paths in 7-week-old mice interfollicular epidermal cell(*15*, *29*). Recently, using also the power of single-cell transcriptomic data, *The Tabula Muris Consortium* assessed cell-type-specific age-related changes in the *Mus musculus* mouse. In skin, they found that the relative abundances of keratinocyte stem cells decreased significantly with age, in contrast with what we found in NMR. Moreover, only 2 of their top 20 differentially expressed genes in skin computed by age (3, 18, 21 and 24 months) were modified through aging in NMR. These 2 genes, *Aqp3* and *Hspa8,* were upregulated in older NMR even if they did not exceed our thresholds and could be of interest for further studies(*44*).

In our study, immune cell cluster 10 was easily distinguished from the other clusters with a characteristic signature *(Cd207, Ctss, Mfge8)* corresponding to Langerhans cells. Interestingly, the level of Langerhans cells appeared unaffected through aging. However, 5 differentially expressed genes, belonging to the *S100* and *Keratin* intermediate filament families, and known to be regulators of inflammation and immunity in skin, were significantly upregulated in older NMR. Again, in mouse and human epidermal aging, there is in contrast a decrease in the number of Langerhans cells. The frequency and maturation of epidermal Langerhans cells have been shown to be reduced in aged mice starting at 12 months of age. Therefore, NMR skin is featured by a maintenance of epidermal stem cell compartment -possibly through *Igfbp3* gene-as well as a maintenance of Langerhans cells density. To date, only one study using sc-RNAseq profiling has focused its attention on NMR immunity where both splenic and circulating immune cells were examined, but not skin. Authors found that NMR immune system was characterized by a high myeloid to lymphoid cell ratio as compared to mice but only healthy young NMR were used(*46*).

It should be noted here that we focused our study on skin epidermal keratinocytes although it has been demonstrated that human dermal fibroblasts undergo a partial loss of their cellular identity during aging(*21*). Of note, alterations in hair follicle stem cell niche -that occur in mice aging-was not possible to address in this glaber rodent(*31*).

We also found in this study that middle-aged NMR showed a histological skin structure similar to the young ones, to the exception that the epidermal thickness was paradoxically increased at higher age as a consequence of increased number of epidermal layers. Marked thickness of the corneous layer in the adult NMR has previously been shown(*10*) and lack of fur has also been described to be compensated by a thicker epidermis(*9*). The reason of such increase remains unknown. It could result from the reaction to the higher cumulative skin friction in their tunnel habitats for older animals. It could also be due to the overexpression of *Igfbp3* in the basal/germinative layer of the epidermis in older animals, leading to a more active stem cells committing to their differentiation pathway in the upper layers.

Alterations in aging have an impact on water loss, body temperature, and wound healing that is delayed, rendering the skin more susceptible to injury. Here, functional experiments of classical surgical excision wounds on the back of NMR showed no significant difference between the 2 age groups, once more in contrast with what has been shown in mouse and human(*14*). For example, wound closure has been proven to occur more rapidly in young mice (6 months) than in mature (15 months, middle-aged) or aged mice (26 months; *47*). Knowing the expression of *Cav1* in epidermal stem cells and its role in healing process and aging(*39*, *40*), the *Cav1* expression found in NMR metacluster basal 1 could explain the maintenance of wound healing despite aging. Therefore, the strong expression of *Cav1* in epidermal stem cells might play a role in regulating proliferative signaling pathways and could offer a clue to the aging resistance observed in NMR.

Immunosenescence i.e. a decrease in immune cells and increased fibroblast senescence is involved in reduced protection against infections and cancers. In the same way, reduction of epidermal thickness and epidermal stem cell compartment is associated with decreased healing abilities and prolonged wounding. Based on their size, NMR should live about 6 years. Therefore, the maintenance of epidermal stem cell and Langerhans cells compartments in NMR -possibly in relation with *Igfbp3* and/or *Cav1* maintained expression-appear to contribute to the remarkable longevity and resistance to aging and diseases of the NMR. Finally, our data also show the potential of the NMR as a model organism for studying cutaneous biology and disease resistance.

## MATERIALS AND METHODS

### Experimental Design

Using single-cell RNA-sequencing (scRNA-seq), histological analysis and wound healing functional tests in young (Y, mean age: 1.1 years) and middle-aged (A, mean age: 11.3 years) naked mole-rats (NMR) epidermis, we aim to better understand the cellular and molecular mechanisms of skin homeostasis and repair in these animals during aging. These results will potentially provide clues to understand why NMR possess a greater lifespan.

### Animals

Y and A males were obtained from four distinct colonies maintained at the Ecole Nationale Vétérinaire de Maisons-Alfort (EnVA). Animal experiments were performed according to experimental protocols following European Community Council guidelines and approved by the Institutional Animal Care and Use Committee of the Ecole Nationale Vétérinaire de Maisons-Alfort (n°016).

### Single-cell RNA sequencing

Epidermal cells from 3 Y and 3 A NMR were used. All the samples were collected and processed at the same time. NMR were anesthetized by the inhalation of 4.9% isoflurane delivered at a flow rate of 300 ml min^-1^ in ambient air and ethanol was applied topically to the dorsal skin for 20 seconds. Punch biopsy devices were used to create 5 mm surgical wounds on the backs of the animals. To preserve cell viability, skin pieces were immediately placed in PBS/BSA 3% medium on ice until dissection. The wounds were rinsed with betadine solution. Using a scalpel, the subcutaneous fat was removed from the dermis. The cleaned tissues were placed dermis side down in a sterile petri dish in a trypsin 0.25%/EDTA 1mM solution at 37°C for 1h. The enzymatic digestion was stopped by addition of PBS/FBS 10%. This floating procedure allows the digestion and dermis-epidermis separation by lifting the epidermis up straight above the dermis. Epidermis was then mechanically dissociated and minced in small pieces with razor blades in PBS and then triturated in 25mL, 10mL then 5mL pipets. 3 rounds of washes were performed starting with 100μm filtration, 70μm and 40μm and interspersed with centrifugation cycles at 150g at 4°C for 5 minutes. Finally, the cell pellet was resuspended in PBS/BSA 3% and cell viability was measured using trypan’s blue method on Kova slides. All single cell suspensions had a viability greater than 75%.

Single cells then were captured in droplet emulsion using the Chromium Controller (10× Genomics, Pleasanton, CA), and scRNA-seq libraries were constructed according to the 10× Genomics protocol using the Chromium Single-Cell 3’ Gel Bead and Library V2 Kit (10× Genomics, Pleasanton, CA). In brief, cell suspensions were diluted in PBS/FBS 3% to a final concentration of 1 × 10^6^ cells/mL (1,000 cells per μL). Cells were loaded in each channel with a target output of 5,000 cells per sample. All reactions were performed in a C1000 Touch Thermal Cycler (Bio-Rad Laboratories, Hercules, CA) with a 96 Deep Well Reaction Module. Twelve cycles were used for cDNA amplification and sample index PCR. Amplified cDNA and final libraries were evaluated using a Bioanalyzer 2100 (Agilent Technologies, Santa Clara, CA) with a high sensitivity chip. Samples were sequenced on an HiSeq 4000 instrument (Illumina, San Diego, CA).

### scRNA-seq pre-processing

scRNA-Seq data analyses were performed by GenoSplice technology (www.genosplice.com). Sequencing data quality analysis was performed using FastQC v0.11.2 on 6 NMR epidermal cells samples, 3 from a Y NMR group (samples 1J, 5J, 6J) and 3 from an A NMR group (samples 2V, 3V, 4V). For read alignment and unique molecular identifiers (UMI) quantification, CellRanger software v3.0.2 was used on Heterocephalus glaber genome HetGla_female_1.0 with default parameters and gene annotation from Ensembl 96. The 6 expression matrices containing the UMI counts were merged, and only the genes with UMI ≥ 1 in at least one cell were kept. The following filters were applied to generate a global matrix used in further analysis: cells with UMI ≥ 2000, number of detected genes ≥ 700, and cells with UMI in mitochondrial genes ≤ 25%. For UMIs normalization, Seurat v3.1.1 was used [PMID:29608179], and global-scaling normalization method was applied with a scale factor of 10,000 and log-transformation of data. This was followed by a scaling linear transformation step, to avoid highly-expressed genes having higher weight in downstream analysis.

### Clustering and marker genes

PCA was performed on the scaled data, with a Jackstraw plot to choose how many PCs to retain as an input for Seurat clustering step. All samples were evenly distributed among clusters, except for sample 2V that showed a very distinctive pattern and was grouped in separated clusters. This sample was therefore removed from the expression matrix in further analysis, keeping 3 samples in the Y NMR group (1J, 5J, 6J) and two samples in the A NMR group (3V, 4V). A second clustering step was performed with the five remaining samples using default parameters, Louvain algorithm as the clustering method and a resolution parameter defining the clusters granularity set to 0.4. Marker genes defining each cluster were found via differential expression testing, with a Wilcoxon rank sum test and a log fold change threshold (logFc) of 1. The found markers were compared to a list of 98 genes of interest for cluster identification. Initial 11 clusters were next grouped in the following metaclusters: Basal1, Basal2, Intermediate! Intermediate2, Corneum and immune cells.

### Trajectory

Single-cell trajectory was constructed using M3Drop [PMID:30590489] and Monocle v2.10.1 [PMID:28825705]. Input genes for Monocle trajectory construction were selected using an unsupervised approach via M3Drop result, which identifies differentially expressed genes based on a Michaelis-Menten function for the relationship between mean expression and dropout rate, the relevant genes being the ones shifting above a fitted curve. The default Monocle workflow was then performed to generate the trajectories, with focus on known markers expression profiles *(Krt14, Krt10, Lor)* to select the appropriate trajectory orientation. We utilized Monocle 2.6.425 to order cells in pseudotime based on their transcriptomic similarity. Pseudotime-dependant genes have been identified using differential GeneTest function from Monocle. Those genes were the most significant in this context and Monocle reconsidered this list of 820 genes and kept the majority of them for the analyses (791 pseudotime-dependent genes). The differential test was performed on genes used to construct Monocle trajectory on cells from Young samples and selected according to q-value (lower than 0.05). Clustering of these genes was done by Monocle functions. Heatmaps for Y and A samples used this clustering of genes.

### Enrichment analysis

Using the Cytoscape v3.7.2 software and BiNGO plugin (binomial statistical test and Bonferroni correction, threshold with adjusted p<0.01), we determined which Gene Ontology (GO) categories are statistically overrepresented in a set of genes of a biological network in the newly defined metaclusters on Mus Musculus annotations. For specific analysis, a Fisher exact-test was performed on a list of genes compiled from REACTOME (ROS) and GO terms (aging and wound healing) databases. Results with uncorrected p-value ≤ 0.05 were considered enriched.

### Histology and immunohistochemistry

Four A and 4 Y NMR skin samples were collected under anesthesia by performing 5mm skin punch biopsies on their back skin. The skin was then immerged in formalin for 12 to 24 hours at 4°C, and embedded either in paraffin or OCT compound for sectioning.

Formalin-fixed paraffin-embedded skin sections (5 μm) were stained with Hematoxylin and Eosin, Sirius Red and Masson’s Trichrome using standard procedures. Epidermal, dermal thickness and number of epidermal layers and cells were measured using

Image J software. Cells were counted in 10^-3^mm^2^ randomized surface squares in the epidermis of each specimen. Quantifications were done on various parts of the skin, dissected in a gridded fashion and 3 different sectors were analyzed; for each different sector of the skin, 3 different areas of 10^-3^mm^2^ were counted.

Formalin-fixed paraffin-embedded skin sections or cryo-sections were labeled with various antibodies (Table S1). Briefly, for formalin-fixed paraffin-embedded sections, a prior antigen retrieval was performed by incubating the slides with citrate buffer (pH=6) or EDTA buffer (pH=9) at 98°C for 20 min. Sections were then washed with Tris buffer saline (TBS) (137mM NaCl and 20mM Tris) / Tween 20 (TBST) 0.1% and blocked for 2 hours with TBST/Normal Goat Serum (NGS) 5%. The sections were incubated with the different primary antibodies diluted in blocking buffer overnight at 4°C for intracellular staining, followed by secondary antibody goat anti-mouse Alexa 546 (dilution 1:1000, Life Technologies), or goat anti-rabbit Alexa 488 (dilution 1:1000, Life Technologies) for 45 min at room temperature. Slides were also incubated with 4’,6-diamidino-2-phenylindole (DAPI, Sigma-Aldrich) for 5 min for nuclei staining and coverslipped using Fluoromount G (Southern Biotech, Birmingham, AL, USA). For hyaluronic acid staining, the slides were incubated 2h with HA-binding protein (at 1,6/100 dilution) with 0.2% bovine serum albumin and 0.02% TritonX-100 in PBS. HA-binding protein staining was achieved with 1/100 strepavidin HRP (Perkin Elmer, Massachusetts, USA) solution for 30 min at room temperature and 3,3’-diaminobenzidine (Liquid DAB + substrate Chromagen System; Dako, Santa Clara, CA, USA) for color development. Finally, sections were counterstained with hematoxylin (Sigma-Aldrich, St. Louis, MO, USA) and mounted using DPX (HA staining in brown). Negative controls were performed in parallel with the samples substituting the primary antibody with the equivalent isotype.

The sections were visualized under an Olympus BX63F microscope equipped with an Olympus DP73 camera (and pictured with Metamorph software) (Olympus, Tokyo, Japan).

### Wound healing procedure

Eight NMR were anaesthetized and punch biopsy devices were used to create four 5 mm surgical wounds on the backs of the animals. All tissues above the panniculus carnosus were excised. Wounds were left uncovered until harvesting. Standardized images of the wounds were obtained at various time points, with a Sony Cybershot 10.1-megapixel DSC-W180 digital camera (Sony, Tokyo, Japan). The images were used to measure the wound closure rate using ImageJ software every 2-3 days.

### Statistics

The data were analyzed using Chi^2^ statistic test, Wilcoxon test and Mann-Whitney U test. n indicates the number of independent experiments performed. In all histograms, asterisks correspond to: *p < 0.05, **p < 0.01, ***p < 0.001, ****p < 0.0001.

## General

We thank Genosplice Technology (www.genosplice.com) for scRNA-Seq data analyses and Daniel Vaiman, Dany Nassar, Bénédicte Oulès, Maria Sbeih and Mathieu Castela for helpful discussions. We also thank Aude Fert, Michèle Oster, Sandrine Lamblin, Romain Morichon (Imaging Platform UMS30 LUMIC), IMAG’IC and HISTIM platforms of Institut Cochin, and Thomas Lilin (Ecole nationale vétérinaire d’Alfort) for their technical support.

## Funding

This work was funded by the Société Française de Dermatologie, Emergence Idex research program-Université de Paris, and Fondation pour la Recherche en Physiologie.

## Competing interests

The authors state no conflict of interest.

## Data and materials availability

Raw and processed scRNA-seq data have been deposited in the GEO database under accession code GSE152886.

## Notes

### Competing Interest Statement

The authors have declared no competing interest.

