## Supplementary Figures 1 to 6 and supplementary Table 1 for "Epidermal stem cell compartment remains unaffected through aging in naked mole-rats"

### Description of Supplementary Materials

#### Supplementary Figures

**Fig. S1**. Quality controls of scRNA-seq.

**Fig. S2**. ScRNA-seq of naked mole-rat epidermal cells define 3 cellular states.

**Fig. S3**. ScRNA-seq of young versus middle-aged naked mole-rat epidermis.

**Fig. S4**. Pseudotime estimation and metacluster identification of young and middle-aged naked mole-rat epidermal cells.

**Fig. S5**. Bubble plot representing the immune Gene Ontology (GO) terms released by the BiNGO analysis.

**Fig. S6**. Epidermal morphology of young and middle-aged naked mole-rats.

#### Supplementary Table S1

Characteristics of antibodies used for immunohistochemistry.

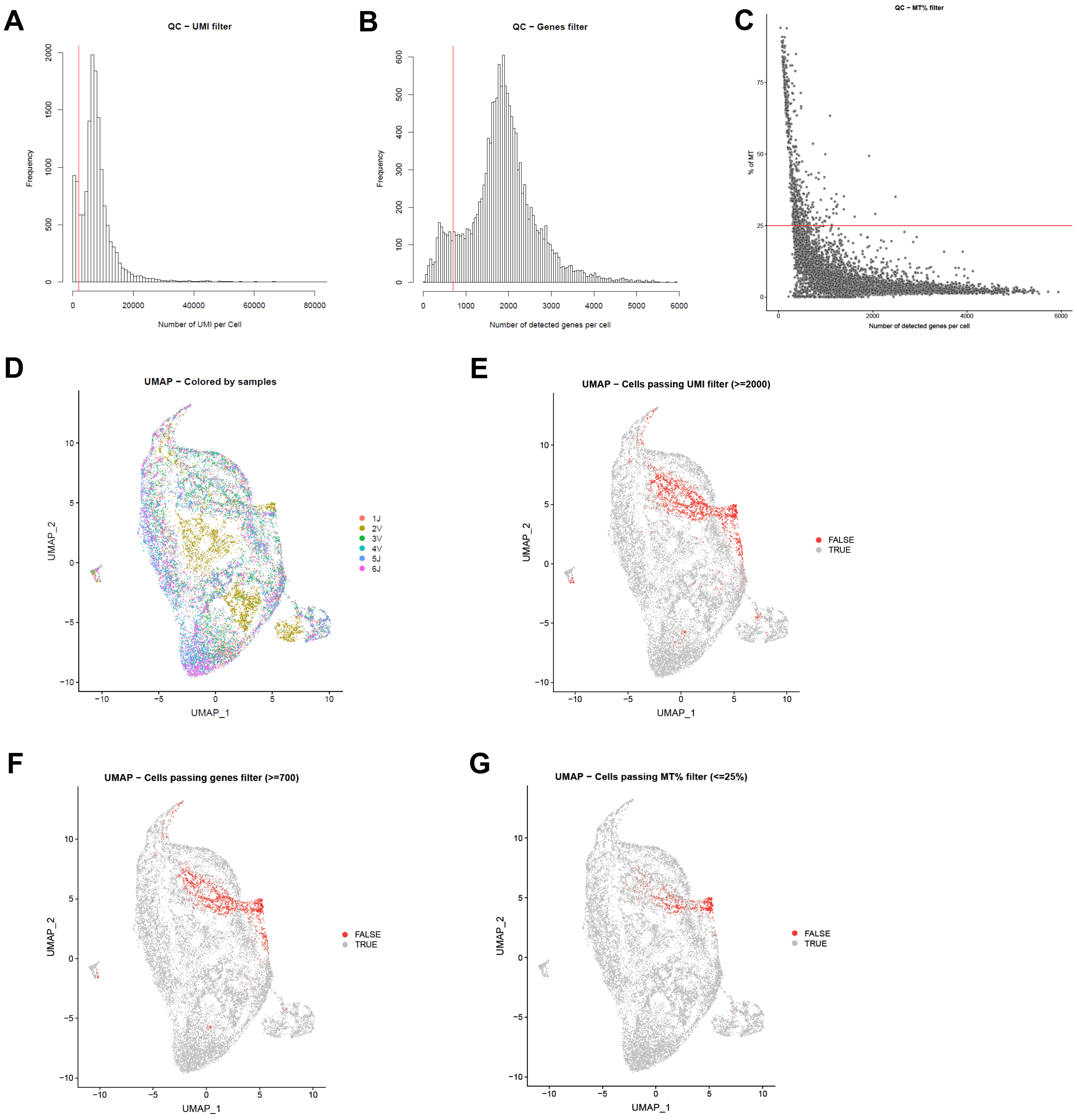


**Fig. S1. Quality controls of scRNA-seq.**
(**A**) Histogram of UMI number per cell. A minimum threshold is applied at 2,000.
(**B**) Histogram of genes count per cell. A minimum threshold is applied at 700.
(**C**) Histogram of percentage of mitochondrial reads per cell. A minimum threshold is applied at 25%.
(**D**) UMAP representation of all cells, colored by sample. Cells from sample 2V are apart from other cells and filtered for further analysis.
(**E, F, G**) UMAP representation of all cells, passing or not each previous QC point (number of UMI per cell, number of genes count per cell and percentage of mitochondrial reads per cell).

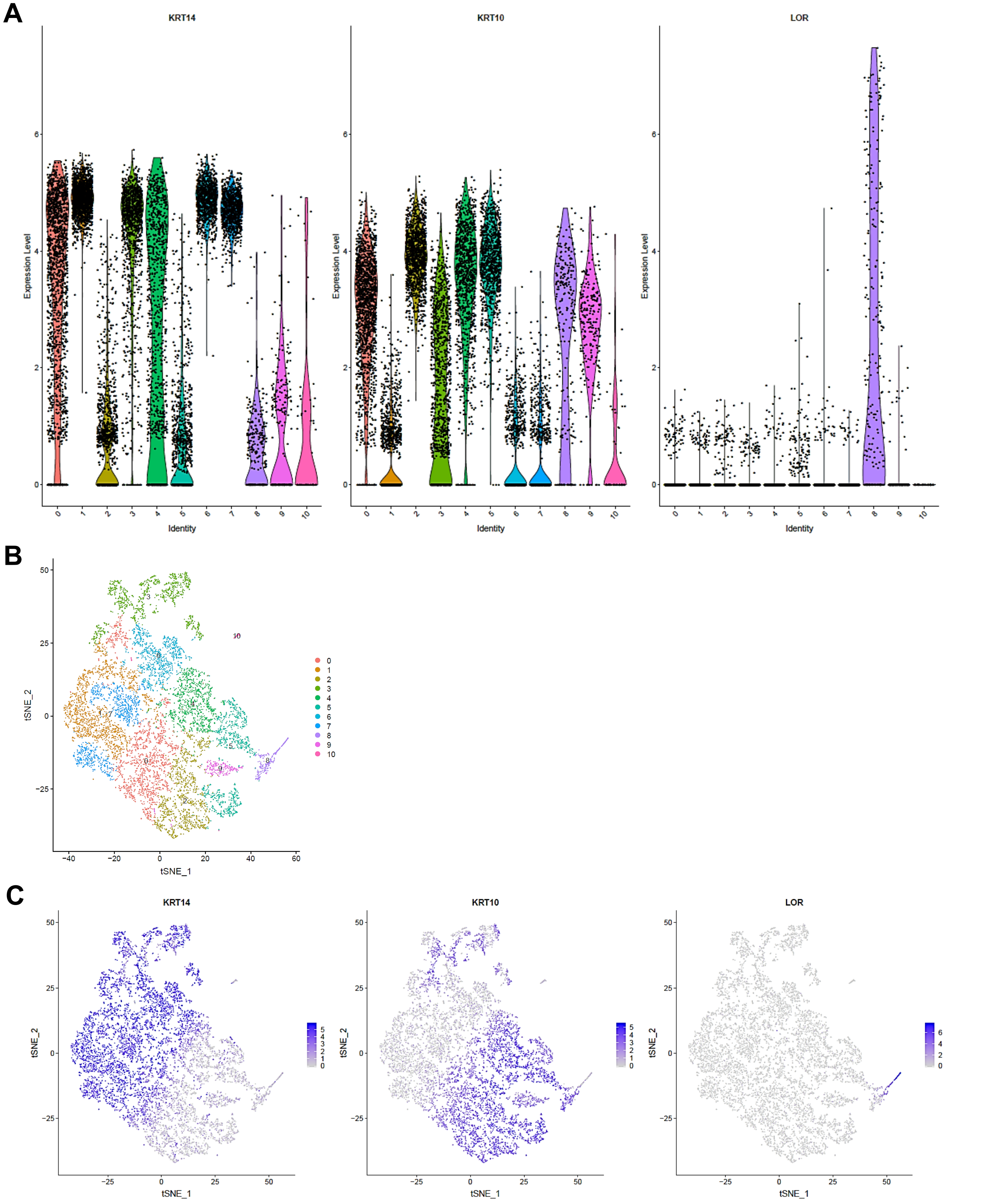


**Fig. S2. ScRNA-seq of naked mole-rat epidermal cells define 3 cellular states.**
(**A**) Violin plots of *Krt14*, *Krt10* and *Lor* marker genes expressed by naked mole-rat (NMR) epidermal cells, leading us to confirm 3 cellular states of keratinization.
(**B**) t-distributed Stochastic Neighbor Embedding (t-SNE) visualization of epidermal cells clustering.
(**C**) t-SNE visualization of *Krt14, Krt10 and Lor* marker genes expressed by NMR epidermal cells.

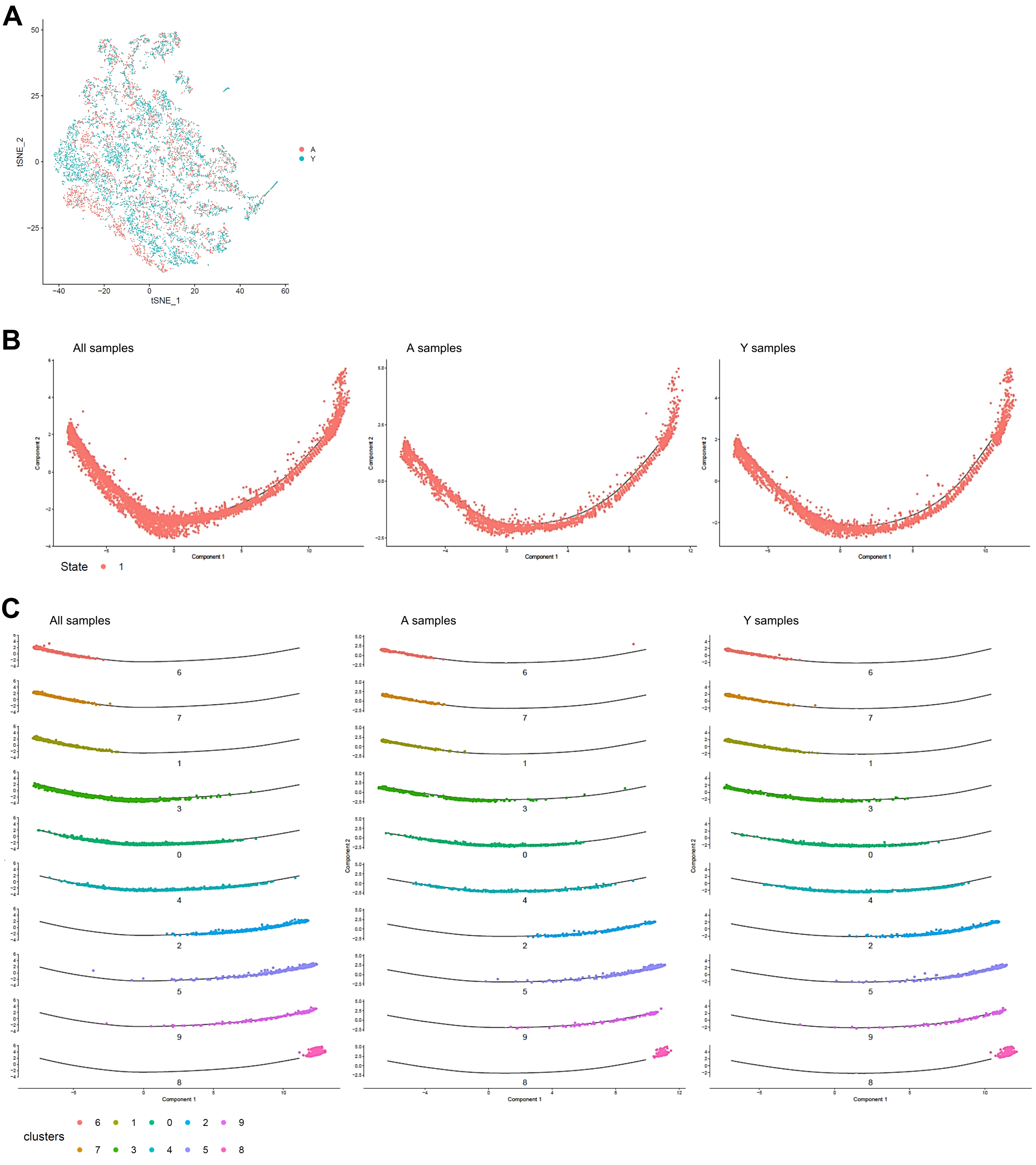


**Fig. S3. ScRNA-seq of young versus middle-aged naked mole-rat epidermis.**
(**A**) t-SNE visualization of epidermal cells where both middle-aged (A, red) and young (Y, blue) naked mole-rats (NMR) were jointly projected on the same plot, showing an overlap of the 2 age groups tested.
(**B**) Unsupervised differentiation trajectories for merged, A and Y keratinocytes colored by Monocle state of differentiation. A single and similar trajectory in both samples was found.
(**C**) Unsupervised differentiation trajectories for merged, A and Y keratinocytes segregated by cluster. No difference between the 2 age groups was found.

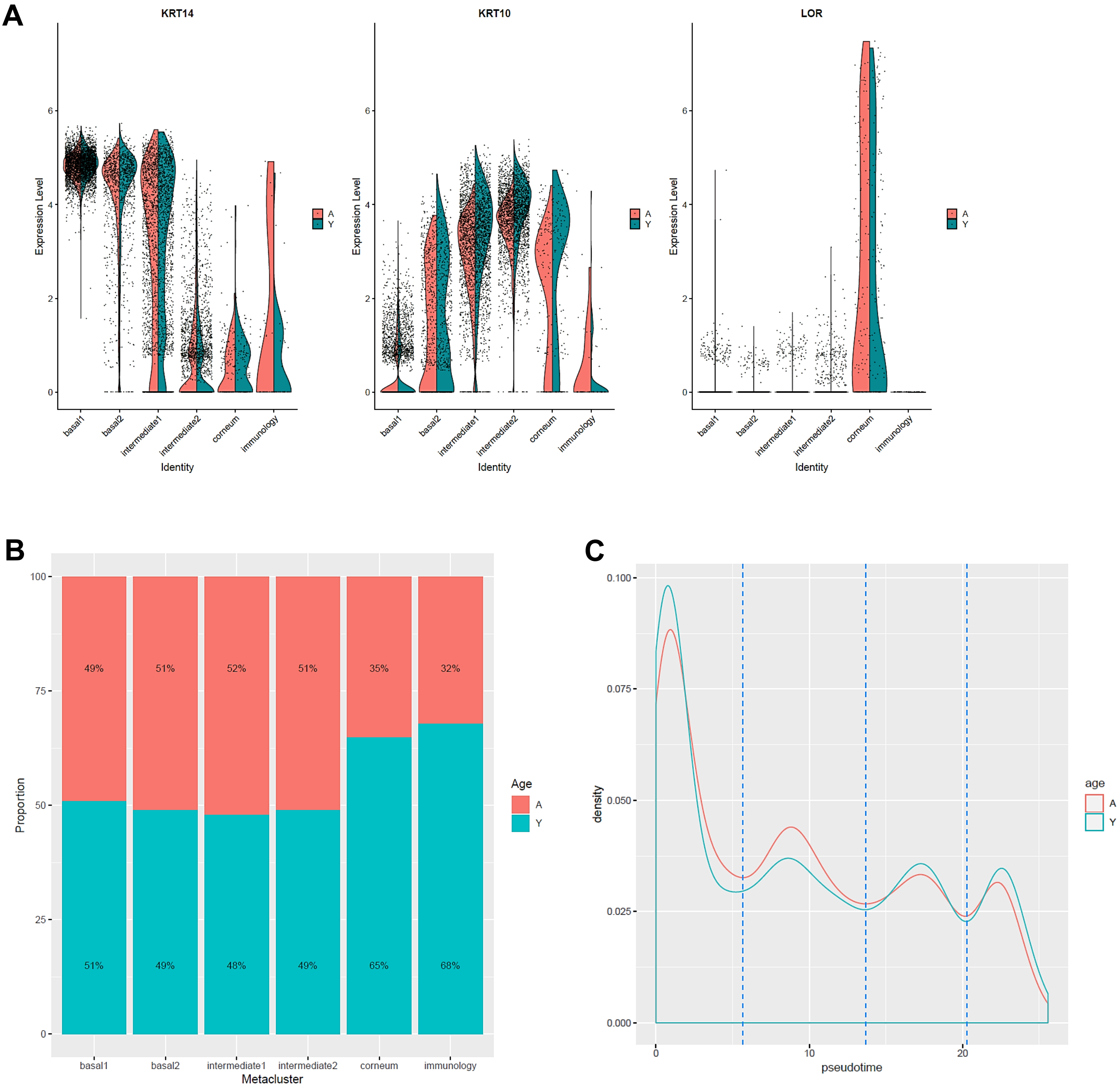


**Fig. S4. Pseudotime estimation and metacluster identification of young and middle-aged naked mole-rat epidermal cells.**

(**A**) The expression of *Krt14, Krt10 and Lor* marker genes visualized with violin plots confirmed the cellular states of the newly defined metaclusters for middle-aged (A, red) and young (Y, blue) naked mole-rats (NMR).

(**B**) Bar graph representing the relative proportion of epidermal cells in each metacluster between A (red) and Y (blue) animals. No significant difference between the 2 age groups was noted (Chi² statistic test).

(**C**) Distributions plotting the frequency of A and Y cells as a function of the pseudotime on its scale. Distributions were similar between the 2 age groups.

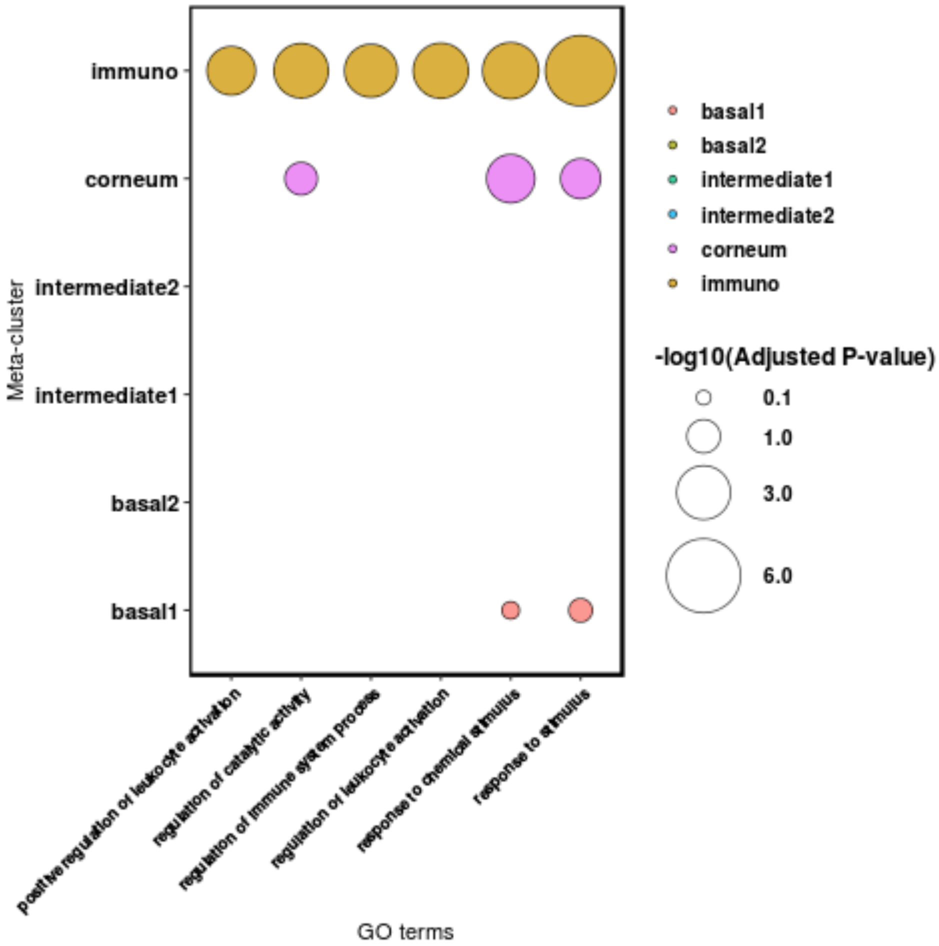


**Fig. S5. Bubble plot representing the immune Gene Ontology (GO) terms released by the BiNGO analysis**. The bubbles represent the adjusted p-value of these terms in the immune cluster and in the other clusters when they emerged.

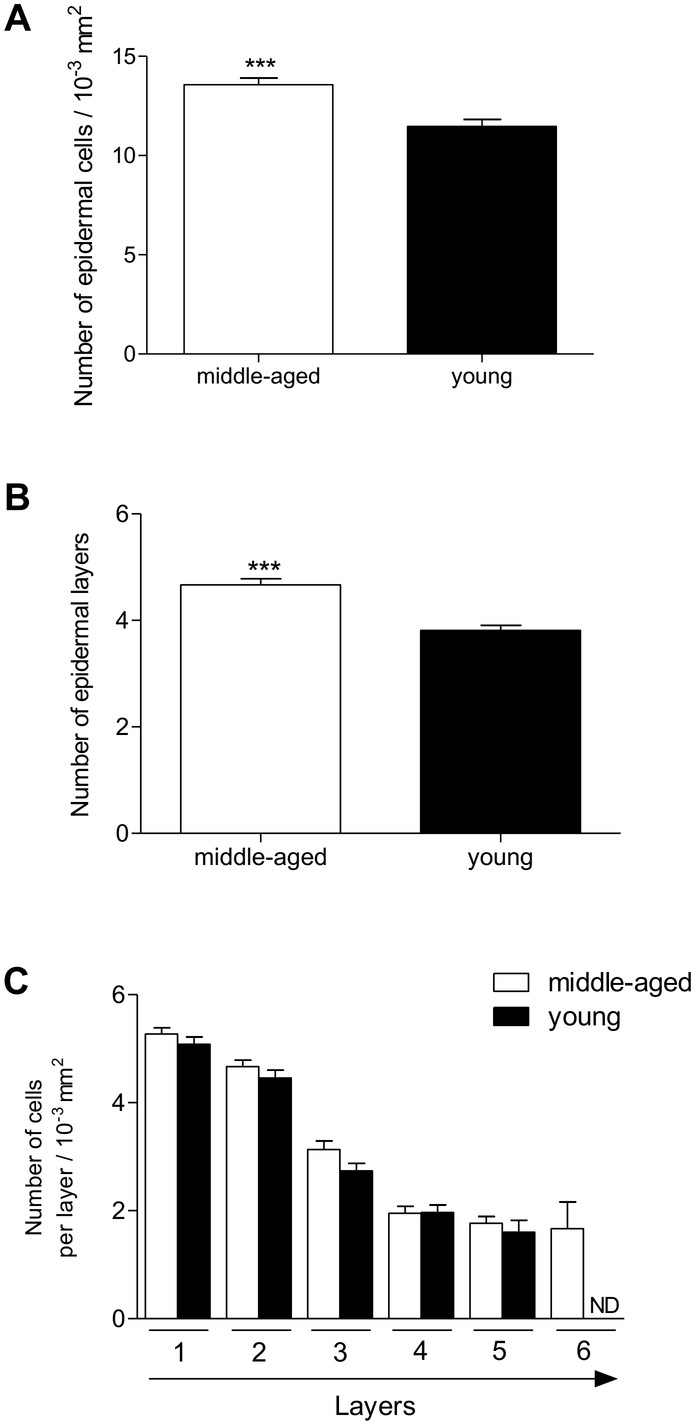


**Fig. S6 : Epidermal morphology of young and middle-aged naked mole-rats.**
(**A**)Histogram showing the number of epidermal cells in middle-aged versus young naked-mole rats (NMR), per surface.
(**B**) Histogram showing the number of epidermal layers in middle-aged vs young NMR.
(**C**) Histogram showing cell number per layer in middle-aged versus young NMR, per surface. An additional external layer was found in aged samples.
n=4 animals per group. Bars: SEM. *represents differences between middle-aged and young animals.***p≤0.0001 in Mann-Whitney U test. ND : non detectable.

**Supplementary Table S1: Characteristics of antibodies used for immunohistochemistry.**

| REAGENT or RESOURCE | DILUTION | SPECIES | SOURCE | IDENTIFIER | SECTION METHOD |
|---|---|---|---|---|---|
| Antibodies | | | | | |
| Keratin 14 | 1:1000 | rabbit | Biolegend | 905301 | paraffin |
| Keratin 10 | 1:1000 | rabbit | Biolegend | 905401 | cryo |
| Loricrin | 1:1000 | rabbit | Biolegend | 905101 | paraffin |
| Filaggrin | 1:500 | rabbit | Biolegend | 905801 | paraffin |
| Laminin 5 | 1:200 | rabbit | Abcam | ab14509 | cryo |
| ITGB4 | 1:250 | rabbit | Abcam | ab182120 | paraffin |
| Ki67 | 1:400 | rabbit | Abcam | ab15580 | cryo |
| Collagen I | 1:250 | rabbit | Abcam | ab21286 | paraffin |
| P16INK4A | 1:100 | rabbit | ThermoScientific | PA1-30670 | cryo |
| Hyaluronic acid | 1,6:100 | bovine | Calbiochem | 385911 | paraffin |
